# The Effect of Eucalyptol on Nursing Home Residents

**DOI:** 10.1101/778209

**Authors:** Seiko Goto, Hinako Suzuki, Toshinori Nakagawa, Kuniyoshi Shimizu

## Abstract

Eucalyptol is one of the most popular volatile components used in many essential oils for relieving sinus and lung congestion caused by variety of conditions. This pilot study aims to analyze clinical evidence of the effect of the scent of eucalyptol on dementia patients to answer two questions: 1) whether eucalyptol aroma is an effectiveness at reducing the symptoms of dementia, and 2) if so, how it mitigates those symptoms. 27 nursing-home residents with dementia were recruited to assess the efficacy of the scent of eucalyptol. Two one-week experiments were performed: the scent was diffused only at wake-up time in the first experiment and at wake-up time and bedtime in the second experiment. Results showed MMSE, DBD, CMAI scores slightly improved in the first experiment and significantly improved in the second experiment even though no subject reported perceiving the scent in either experiment. The present results indicate that eucalyptol is effective in mitigating dementia symptoms in an elder population with an impaired sense of smell.

## I. Introduction

Aromatherapy with essential oils has been used in many countries from ancient times. Aromas trigger physical and mood improvement via the olfactory system.^1,2^Inhaled air containing a scent can reach the circulatory system but also stimulate brain areas directly via the olfactory epithelium. Once the signals reach the olfactory cortex, release of neurotransmitters—serotonin, for example—takes place, which results in the expected effect on emotions.^3,4,5^ Previous studies by Goto showed that visiting a small indoor garden with 20 pots of chrysanthemums for 15 minutes improved the mood of elder subjects with dementia. ^6^ Subjects became more alert and their mood significantly improved while observing the garden. Because some subjects remarked about the floral scent, the effects of visiting the garden were hypothesized to involve olfactory stimulation.^7,8^

However, different parts of plants emit different scents, and scent from the same parts can differ based on freshness. Even within the same plant family, different varieties can have very different volatile components.^9^ Because the volatile components of plants are uncertain, components of volatile organic compounds in commercial essential oils bearing the same plant name, i.e. lavender or rose, differ by brands. Furthermore, although essential oils are composed of multiple volatile components, there has been little in-depth research on the efficacy of a single component. Although inhalation of essential oils has been traditionally practiced as aromatherapy, the concentrations, dosage, duration, and frequency of usage are not standardized in practice.^10^

Because the mode of administration of an essential oil and its content, as well as the effects of each volatile, are not clear, the great majority of aromatherapy is of scientifically inadequate quality.^11^ Even if the scent is usually effective, using different essential oils with different volatile components at different concentrations and frequencies may not have the desired effect, or cause problems. It is important to identify the effects of a single volatile with controlled dosage and duration, particularly if the scent is used for a frail population, such as elderly people with dementia. Therefore, this study focused on eucalyptol, one of the main volatiles of chrysanthemums,^12,13^ measuring the effects of its scent on elderly Japanese adults with dementia in a nursing home.

## II. Methods

### 1. Study Design

Eucalyptol (1,8 cineole) is a monoterpenoid compound present in large amounts in a variety of plants often used in cosmetics.^14,15^ Eucalyptol has been used as an ingredient in many medicinal products to treat bronchitis, sinusitis, and chronic rhinitis, and also asthma, and rhinosinusitis.^16,17^ Eucalyptol compound also enhances blood circulation,^18^ and Ambrosch reported that dermal application of eucalyptol increased brain activation in the frontal cortex and significantly improved brain function related to working memory in rats.^19^

Two one-week single blind experiments were conducted between 6/18-6/24/2018, and 10/12-10/18/2018, approximately 4 months apart. Before and after each test, subjects and controls were tested using the MMSE (Mini-Mental State Examination), DBD (Dementia Behavior Disturbance scale), and CMAI (Cohen-Mansfield Agitation Inventory) (6/8-6/17, 6/25-7/3, 10/2-10/11, 10/19-10/25). During the second experiment, caregivers gave a simple memory test to the subject group. Follow-up DBD, CMAI, and MMSE tests were conducted for the control group three weeks after the second experiment (11/6).

In the first experiment, 8 drops of eucalyptol oil were diffused without dilution with an electric diffuser (Truffle) produced by the Tree of Life Company (Japan) set on a side table approximately 50 cm away from the subject’s pillow, and started one hour before the subject’s wake time, and run for 60 min. In the second experiment, the same amount of eucalyptol was diffused in the same setting for one hour before the wake time and also for 60 minutes at bed time. Except for diffusing the scent in the subjects’ rooms, no change was made in their daily routine.

### 2. The site

The experiment site is located in Nozomi-no-mori (N. House) in Nagasaki, Japan. N. House is a facility providing daycare as well as short- and long-term stays for elders needing care. The long-term stay facility is comprised of a group home for elders with mild disabilities and a nursing home for those with serious disabilities. There are 9 single rooms in the group home, and 5 units of 10 single rooms in the nursing home. Residents are assigned to either the group home or nursing home based on their Care Level, a level assessed by seven scales established by the Ministry of Health and Welfare in Japan based on needed time of care per day: Support Level 1 and 2, and Care Level 1 to 5.

N. home assigns people from Support Level and Care Need Level 1 and 2 to the group home, and people with Care Need Levels 3 and above to the nursing home. 7-10 staff members are assigned to each unit with 2-3 for 3 work shifts. The policy of the N. Home is to care for residents according to their accustomed lifestyle. Therefore, the N. Home does not set times for waking, eating, bathing, toileting, and going to bed. They serve meals when residents desire them. Although they provide cultural activities such as crafts, calligraphy, and singing in each unit twice a week, they do not force residents to participate. Since the nursing home does not have a medical doctor, it does not provide medication. Residents either visit the nearby hospital or call a doctor if necessary.

The group home is composed of 9 single rooms (5mx3m) with a common kitchen and living space. The nursing home is composed of 5 units of 10 (4mx3m) single rooms with a common living room. Each room has a bed, a side table, a closet, and chest, and the layout of the room is identical. The temperature is controlled by the air conditioners installed on the ceiling of each room and the windows are closed most of the day. Fig.1 shows the floor plans of the group home and the nursing home (unit 1 and unit 2). Blue indicates rooms of the subject group and pink indicates rooms of the control group. One subject moved from Group Home to the nursing home between 1^st^ and 2^nd^ experiment because of the decline of Care Level.

**Fig 1:**
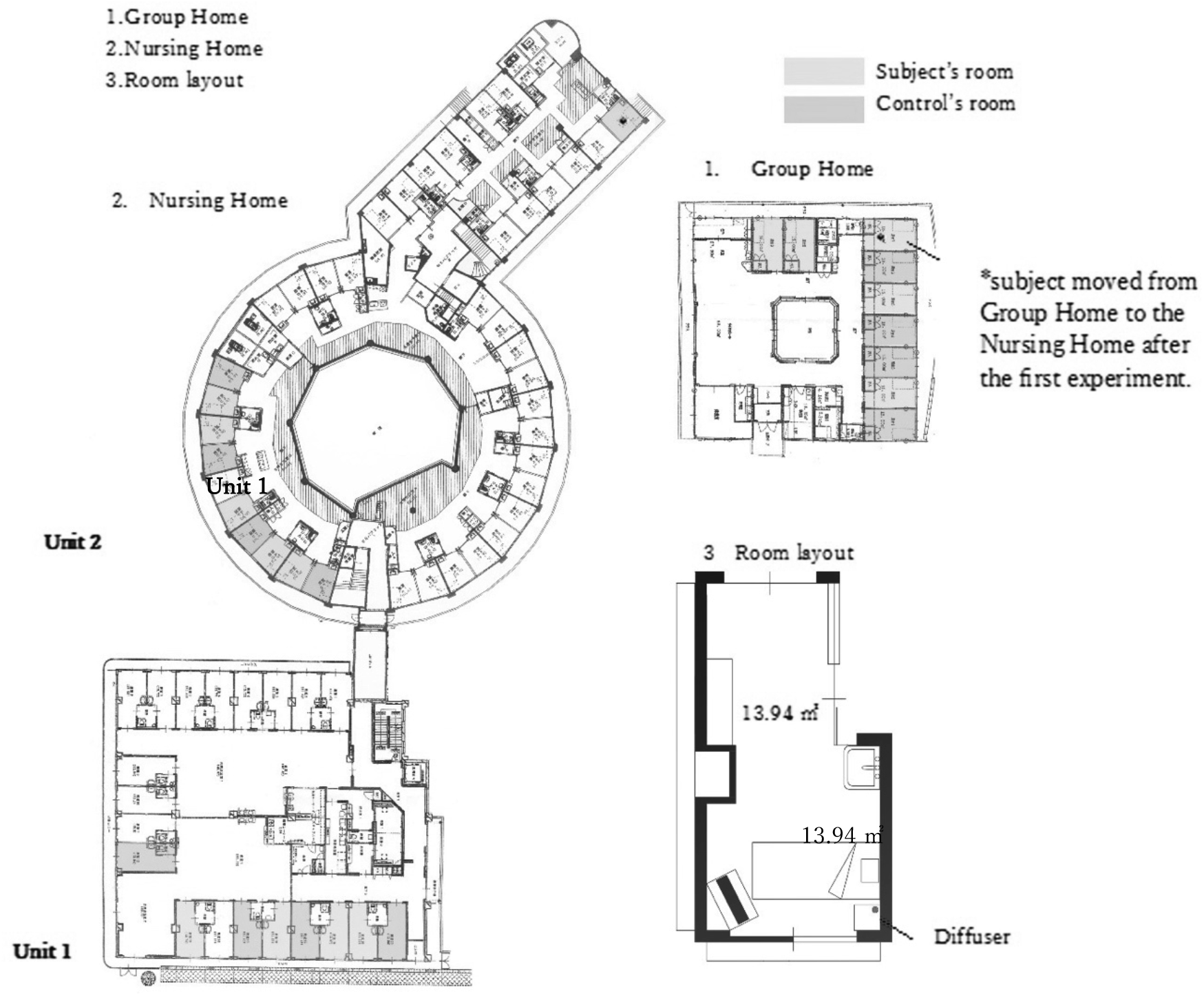
The floor plan of the group home and the nursing home, and the typical layout of the residents’ room.

### 3. The diffuser and the level of the scent

Eucalyptol was diffused by a Truffle diffuser, which is a type of nebulizing diffuser which does not use water and produces a potent mist. It operates on a 2-minute-on and 1-minute-off interval cycle for 2 hours. One Truffle is designed to fill up to a 32m^2^ room with scent in 2 minutes. The temperature of the experiment rooms was maintained at 22 ± 1℃. The diffuser was located 50 cm away from the pillow. After 10 minutes of the operation of the diffuser, the scent was noticeable to the experimenters in the room. The level of eucalyptol in the air at the pillow, 50 cm away from the diffuser, was 15.56±1.16 μg/m^3^ in Group Home, 17.04±2.77 μg/m^3^ in Unit 1, and 20.17±3.51 μg/m^3^ in Unit 2. The level of eucalyptol was calculated by averaging the results of 3 air samples collected for 1 minute using a TENAX TA tube (GERSTEL GmbH Co.KG) set in the pump with the flow rate of 0.05 ℓ/min. after 15 minutes of operation of the diffuser.

### 4. Subjects

Experiments were performed in accordance with relevant guidelines and regulations of Nagasaki University with agreement of N Home. Human research protocols for the study were approved by the Ethics Committee of Nagasaki University. The Inclusion and Exclusion Criterion for Study Selection was sufficient health stability to be able to live in the unit during the entire study period. Subjects for the experiment were recruited from the group home and 2 units of the nursing home of N. House (Unit 1 and Unit 2), where residents’ conditions were relatively stable according to N. House’s records. 27 subjects’ family members gave written informed consent.

The 27 participants in the study were divided into two groups: 14 subjects—11 female (average age 83.3 ±8.2) and 3 male (average age 86.3 ±8.0)—and 13 controls—10 female (average age 89.8 ±4.5) and 2 male (average age 81.5 ±0.7). No participants were taking any medication or seeing doctors during the period of experiments. The level of dementia of N. Home residents is assessed by the expert staff using the seven scales of Level of Independent Living established by the Ministry of Health and Welfare in Japan: I, IIa, IIb, IIIa, IIIb, IV, and M. (Medical).

Assignment of Level of Independence is based on the severity of symptoms of dementia as follows:

Level I: able to live independently.

Level II: unable to live independently without assistance with symptoms such as getting lost, and being unable to shop, pay bills, and receive phone calls.

Level III: needing assistance with daily routines, such as eating and bathing, and for dealing with behavior such as screaming and wandering.

Level IV: needing full-time care for dementia with other psychiatric symptoms. Level M: needing medication.

According to the residents’ record of N. Home, the Level of Independence of both groups was classified as follows (the number in parenthesis is the number of residents):

Subject group: IIa (2), IIb (1), IIIb (8), IV (3)

Control group: IIb (2), IIIa (5), IIIb (4), IV (2)

### 5. Questionnaire

Before and after the two experiments, subjects and controls were tested for their level of cognitive impairment and behavioral symptoms using MMSE, DBD and CMAI. Furthermore, a simple memory test was given to the subjects every day by caregivers during the second experiment to test for improvement in recognition of the time and short-term memory. In order to confirm the effect of the scent, follow-up tests with the control group of DBD and CMAI tests, and the memory test with the subject group were conducted 3 weeks after the 2^nd^ test.

#### MMSE

The Mini–Mental State Examination (MMSE) is a 30-point questionnaire to measure cognitive impairment. The test takes between 5 and 10 minutes and examines functions such as attention, calculation, recall, language, ability to follow simple commands, and orientation.

#### DBD and CMAI scale

The Dementia Behavior Disturbance Scale (DBD scale) is a questionnaire for caregivers to measure the behavioral symptoms of patients. The DBD scale is comprised of the following 28 questions about dementia-related behavior, scored by an observer on a 5-point scale (Score 0: Never; 4: Have the symptom all the time).

1. Asks the same question over and over again.
2. Loses, misplaces, or hides things.
3. Shows lack of interest in daily activities.
4. Wakes up at night for no obvious reason.
5. Makes unwarranted accusations.
6. Sleeps excessively during the day.
7. Wanders aimlessly outside or in the house during the day.
8. Repeats the same action (e.g., wiping table) over and over again.
9. Is verbally abusive, curses.
10. Dresses inappropriately.
11. Screams for no reason.
12. Refuses to be helped with personal care tasks, such as bathing, dressing, brushing teeth.
13. Hoards things for no obvious reason.
14. Moves arms or legs in a restless or agitated way.
15. Empties drawers or closets.
16. Wanders in the house at night.
17. Gets lost outside.
18. Refuses to eat.
19. Overeats.
20. Is incontinent of urine (wets himself/herself).
21. Paces up and down.
22. Makes physical attacks (hits, bites, scratches, kicks, spits.)
23. Cries or laughs inappropriately.
24. Engages in inappropriate behaviour.
25. Exposes himself/herself indecently.
26. Destroys property or clothing, breaks things.
27. Is incontinent of stool (soils himself/ herself).
28. Throws food.

Cohen-Mansfield Agitation Inventory (CMAI) is a questionnaire for caregivers to assess agitation related to cognitive impairment. The questionnaire consists of the following 14 questions relating to aggressive behaviors and 15 relating to non-aggressive behaviors on a 7-point scale.

##### Physical / Aggressive

1. Hitting (including self)
2. Kicking
3. Grabbing people
4. Pushing
5. Throwing things
6. Biting
7. Scratching
8. Spitting
9. Hurting self or others
10. Tearing things or destroying property
11. Making physical sexual advances

##### Physical / Non-Aggressive

12. Pacing, aimless wandering
13. Inappropriate dressing or disrobing
14. Trying to get to a different place
15. Intentional falling
16. Eating / drinking inappropriate substances
17. Handling things inappropriately
18. Hiding things
19. Hoarding things
20. Performing repetitive mannerisms
21. General restlessness

##### Verbal / Aggressive

22. Screaming
23. Making verbal sexual advances
24. Cursing or verbal aggression

##### Verbal / Non-aggressive

25. Repetitive sentences or questions
26. Strange noises (weird laughter or crying)
27. Complaining
28. Negativism
29. Constant unwarranted requests for attention or help

#### Memory Test

In the second experiment, a simple memory test was given to the subject group in the morning by caregivers. The test consisted of 2 questions: 1. A question about time: “What day is today?”, and 2. A question about short-term memory using cards. For the short-term memory test, the caregiver first showed the subject two cards of fruit and then showed a card with six fruits and asked which fruits the subject saw previously in the cards.

**Fig 2:**
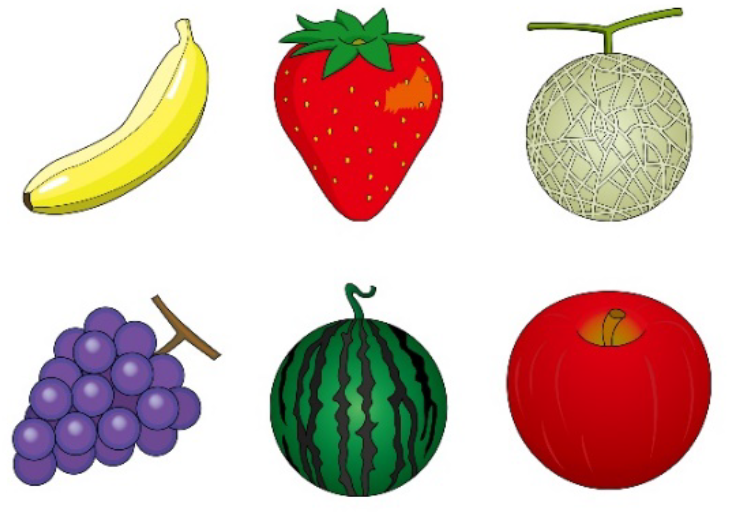
Memory test sample

Significance was analyzed using t-test and established at *p*< 0. 05: * and the trend was established at *p*< 0. 2. For the analysis of change in MMSE score, three subjects whose initial MMSE = 0 were excluded. To assess the interaction among dependent variables, stepwise regression methods were performed for the change in MMSE, CMAI, and DBD using the subject or control, subjectsʼ age, gender, the Level of Care Level, and the Level of Independence as dependent variables.

## III. Results

During the two one-week experiments, two subjects died. All residents followed their routine schedule in their unit, and no subject dropped out, or participated in a new activity during the experiment weeks.

### 1. MMSE

Although the scent of eucalyptol was noticeable to all caregivers and staff members, no participants in the subject and control group noticed it. However, although the change was not significant, the average MMSE of both the subject and control group before and after the first experiment improved. After improving MMSE during the 1^st^ test, the average MMSE of the subject group decreased 2.3 points over the approximately 4 months between the two experiment periods, which means that the dementia advanced while the subjects were not exposed to the eucalyptol scent. However, in the second experiment, after the scent was diffused in the morning and the evening in the subjects’ room for one week, scores of the subject group significantly improved from 7.2 to 9.5 (p=0.002). On the other hand, the average MMSE of the control group remained at the same level. (Figure 3) A stepwise regression showed that diffusing the scent in the room was the possible predictor (p=0.068) in the first experiment and the significant predictor (p=0.018) in the second experiment.

(Fig 3: Result of MMSE)

### 2. DBD and CMAI

Higher scores on the DBD and CMAI indicate worse behavior, so a negative number from before score − after score means improvement in behavior after one week of the experiment. Both subjects and the control group improved their scores in both DBD and CMAI in the second experiment, and the control group improved in Non-Aggressive CMAI even in the first experiment (Fig.4–6).

**Fig 4:**
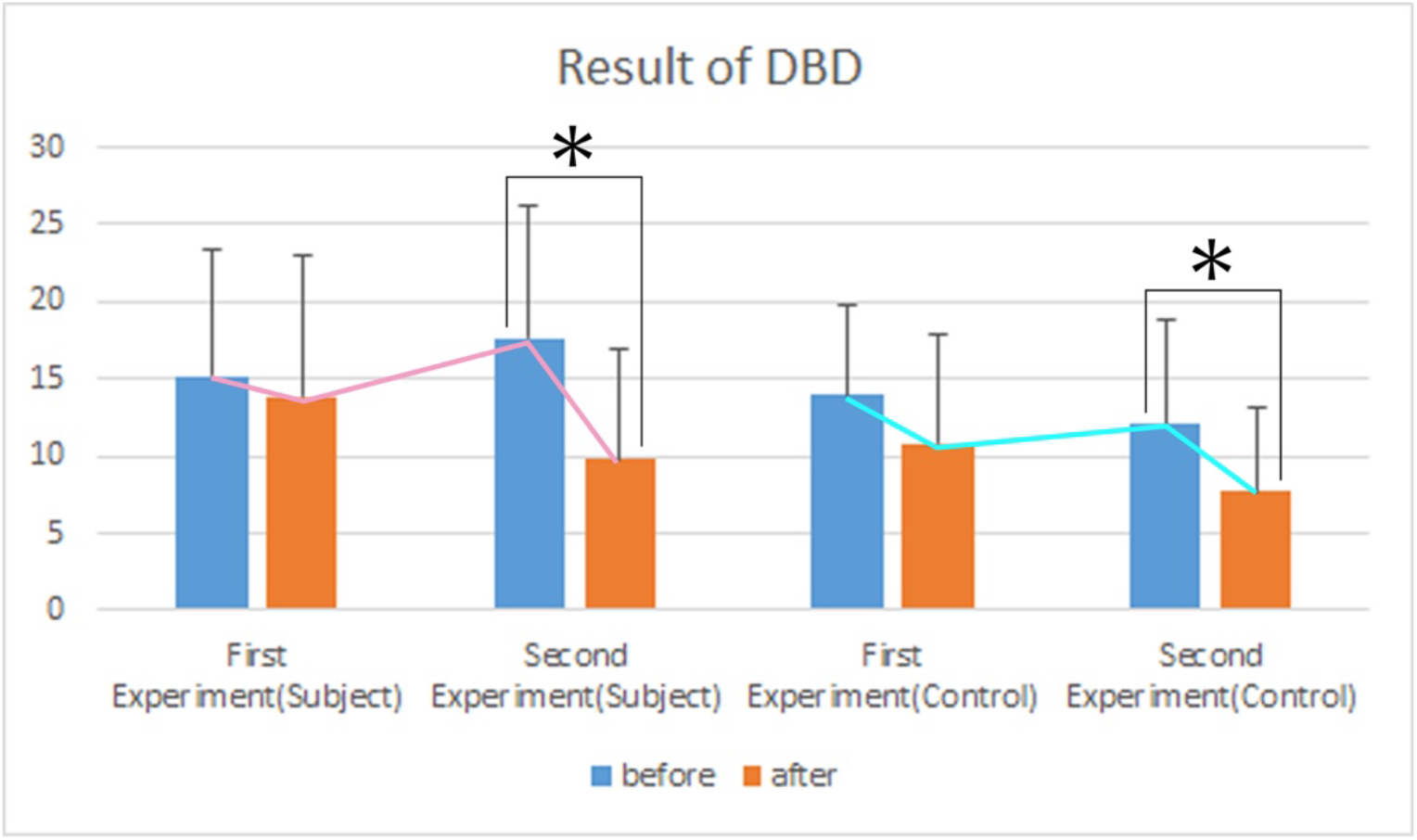
Result of DBD

**Fig 5:**
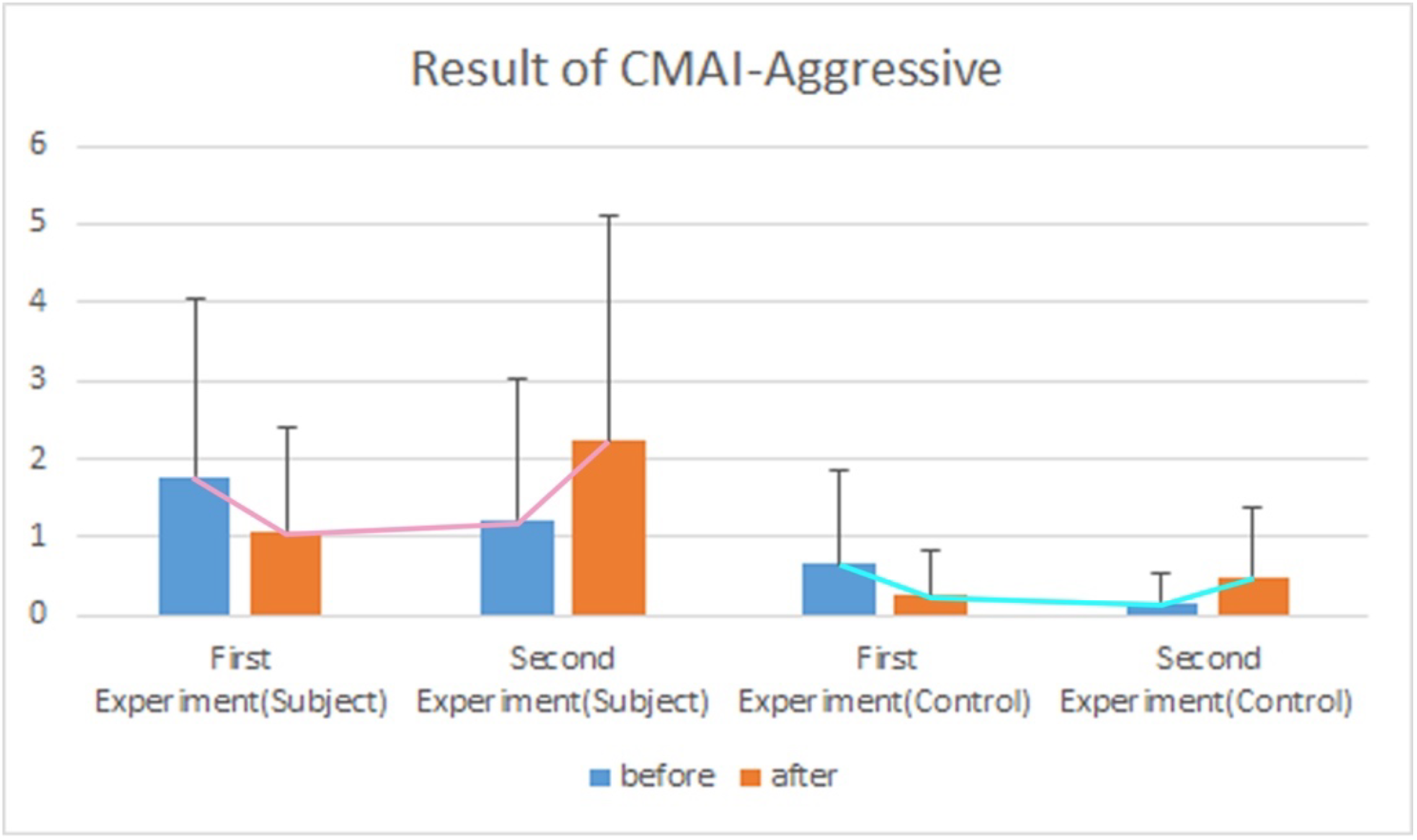
Result of CMAI-Aggressive

**Fig 6:**
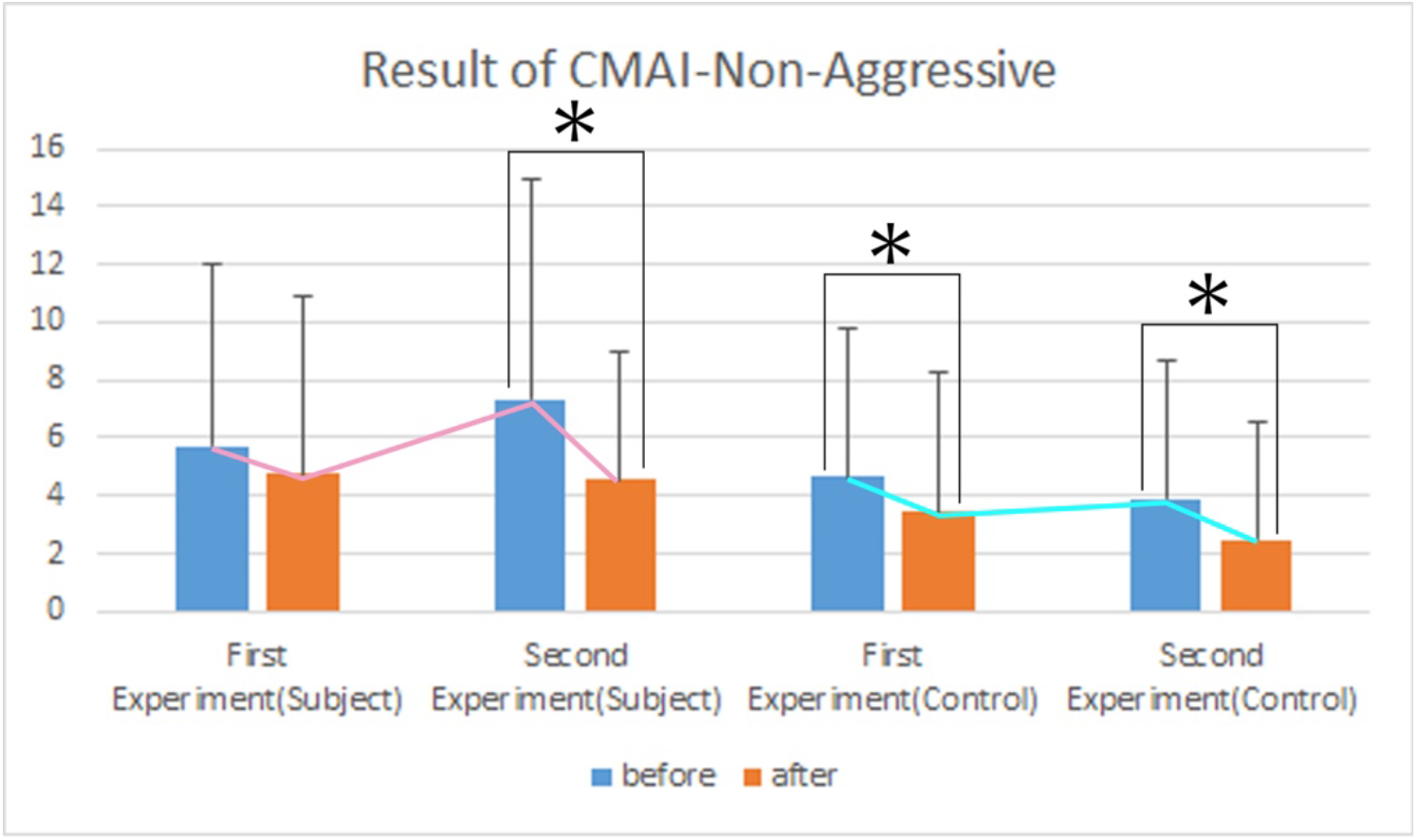
Result of CMAI-Non-Aggressive

Follow-up tests with the control group of DBD and CMAI tests conducted 3 weeks after the 2^nd^ test did not show any improvement in scores, which confirmed that there had been an effect of overflow scent during the experiment week: perhaps the control group was affected by the drifting eucalyptol scent in the living room where they spend the day. Whereas DBD and Non-Aggressive CMAI scores of both the subject and control groups declined during the period of the two experiments, the CMAI aggressive score of both groups stayed almost the same.

The results of a stepwise regression for DBD and CMAI differed from the MMSE. For DBD, it was not whether or not the scent was diffused in the room, but the Level of Independence that was the possible predictor (p=0.1) in the first experiment and the significant predictor (p=0.042) in the second experiment. This means that subjects with a higher Level of Independence had an improved DBD even without the scent being diffused in their room. By contrast, for the CMAI-aggressive, gender and age were significant predictors in the first (gender: p=0.02, age: p=0.007) and the second experiment (gender: p=0.059, age: p=0.018), which means that the aggressive behavior of elder female subjects diminished more.

Table 1–4 shows DBD and CMAI categories which showed a trend and significant change before and after the experiment. Red indicates significant change, orange indicates trend, and grey indicates the categories changed for worse. Although there were categories which showed the decline of behavior, most trends and significant changes showed improvement of behavior. Within the categories of DBD and CMAI, the subject group showed a trend of improvement in 7 categories of DBD in the first experiment. For Non-Aggressive CMAI, 5 categories were improved: Hurting self or others (p=0.19), Eating inappropriate substances (p=0.16), Hoarding things (p=0.16), Cursing or verbal aggression (p= 0.16), Repetitive sentences (p=0.05) and Constant unwarranted requests (p= 0.19).

**Table 1:** DBD categories with trend and significant improvement in the First experiment A: Subject, B: Control

| DBD | First Experiment : Subject |  |  |
| --- | --- | --- | --- |
|  | before | after | p value |
| Overeats. | $0.7 \pm 1.1$ | $0.5 \pm 0.8$ | 0.08 |
| Destroys property or clothing, breaks things. | $0.0 \pm 0.0$ | $0.3 \pm 0.6$ | 0.1 |
| Dresses inappropriately. | $0.9 \pm 1.3$ | $0.3 \pm 0.6$ | 0.15 |
| Repeats the same action (e.g., wiping table) over and over again. | $0.5 \pm 0.8$ | $0.3 \pm 0.6$ | 0.17 |
| Is verbally abusive, curses. | $0.0 \pm 0.0$ | $0.2 \pm 0.4$ | 0.17 |
| Refuses to be helped with personal care tasks, such as bathing, dressing, brushing teeth. | $0.7 \pm 1.1$ | $0.5 \pm 0.9$ | 0.19 |
| Gets lost outside. | $0.5 \pm 1.0$ | $0.2 \pm 0.6$ | 0.19 |

| DBD | First Experiment : Control |  |  |
| --- | --- | --- | --- |
|  | before | after | p value |
| Loses, misplaces, or hides things. | $1.1 \pm 1.2$ | $0.7 \pm 0.9$ | 0.02 |
| Shows lack of interest in daily activities. | $1.8 \pm 1.5$ | $1.3 \pm 1.5$ | 0.05 |
| Sleeps excessively during the day. | $1.4 \pm 1.5$ | $1.0 \pm 1.4$ | 0.14 |
| Makes unwarranted accusations. | $0.4 \pm 0.8$ | $0.1 \pm 0.3$ | 0.17 |
| Repeats the same action (e.g., wiping table) over and over again. | $0.6 \pm 1.0$ | $0.4 \pm 1.0$ | 0.17 |
| Is verbally abusive, curses. | $0.8 \pm 1.3$ | $0.2 \pm 0.6$ | 0.17 |
| Empties drawers or closets. | $0.1 \pm 0.3$ | $0.3 \pm 0.5$ | 0.17 |
| Is incontinent of stool (soils himself/ herself). | $1.5 \pm 1.2$ | $1.3 \pm 1.2$ | 0.17 |
| Wakes up at night for no obvious reason. | $0.7 \pm 1.0$ | $0.4 \pm 0.9$ | 0.19 |

**Table 2:** DBD categories with trend and significant improvement in the Second experiment A: Subject, B: Control

| DBD | Second Experiment : Subject |  |  |
| --- | --- | --- | --- |
|  | before | after | p value |
| Shows lack of interest in daily activities. | 2.4 ± 1.1 | 1.0 ± 0.9 | 0.0004 |
| Sleeps excessively during the day. | 1.6 ± 1.0 | 0.9 ± 1.0 | 0.002 |
| Is incontinent of stool (soils himself/ herself). | 1.8 ± 1.5 | 0.5 ± 1.0 | 0.009 |
| Is incontinent of urine (wets himself/herself). | 1.9 ± 1.4 | 0.7 ± 1.0 | 0.009 |
| Overeats. | 0.5 ± 0.7 | 0.1 ± 0.3 | 0.02 |
| Screams for no reason. | 0.5 ± 0.9 | 0.2 ± 0.4 | 0.05 |
| Asks the same question over and over again. | 1.2 ± 1.3 | 0.9 ± 1.3 | 0.08 |
| Wakes up at night for no obvious reason. | 1.0 ± 0.9 | 0.5 ± 0.8 | 0.08 |
| Repeats the same action (e.g., wiping table) over and over again. | 0.5 ± 0.8 | 0.2 ± 0.4 | 0.1 |
| Refuses to eat. | 0.6 ± 1.0 | 0.2 ± 0.4 | 0.11 |
| Paces up and down. | 0.5 ± 0.8 | 0.3 ± 0.8 | 0.17 |
| Wanders in the house at night. | 0.2 ± 0.6 | 0.1 ± 0.3 | 0.17 |

| DBD | Second Experiment : Control |  |  |
| --- | --- | --- | --- |
|  | before | after | p value |
| Asks the same question over and over again. | 1.3 ± 1.3 | 1.0 ± 1.3 | 0.04 |
| Gets lost outside. | 0.6 ± 0.9 | 0.0 ± 0.0 | 0.05 |
| Makes unwarranted accusations. | 0.4 ± 0.7 | 0.2 ± 0.4 | 0.08 |
| Is verbally abusive, curses. | 0.3 ± 0.5 | 0.1 ± 0.3 | 0.08 |
| Hoards things for no obvious reason. | 0.3 ± 0.5 | 0.1 ± 0.3 | 0.08 |
| Sleeps excessively during the day. | 1.1 ± 1.2 | 0.5 ± 0.8 | 0.09 |
| Is incontinent of urine (wets himself/herself). | 1.6 ± 1.2 | 1.0 ± 0.7 | 0.11 |
| Loses, misplaces, or hides things. | 0.5 ± 0.7 | 0.8 ± 0.8 | 0.17 |
| Dresses inappropriately. | 0.4 ± 0.8 | 0.3 ± 0.6 | 0.17 |
| Paces up and down. | 0.5 ± 1.0 | 0.3 ± 0.9 | 0.17 |
| Is incontinent of stool (soils himself/ herself). | 1.2 ± 1.4 | 0.8 ± 0.9 | 0.18 |

**Table 3:** CMAI categories with trend and significant improvement in the First experiment A: Subject, B: Control

| CMAI | First Experiment : Subject |  |  |
| --- | --- | --- | --- |
|  | before | after | p value |
| Repetitive sentences or questions | 1.1 ± 1.3 | 0.7 ± 1.2 | 0.05 |
| Eating / drinking inappropriate substances | 0.0 ± 0.0 | 0.2 ± 0.4 | 0.17 |
| Cursing or verbal aggression | 0.2 ± 0.6 | 0.1 ± 0.3 | 0.17 |
| Hurting self or others | 0.2 ± 0.6 | 0.0 ± 0.0 | 0.19 |
| Hoarding things | 0.9 ± 1.3 | 0.6 ± 0.9 | 0.19 |
| Constant unwarranted request for attention or help | 0.1 ± 0.3 | 0.3 ± 0.6 | 0.19 |

| CMAI | First Experiment : Control |  |  |
| --- | --- | --- | --- |
|  | before | after | p value |
| Repetitive sentences or questions | 1.0 ± 1.4 | 0.3 ± 0.9 | 0.09 |
| General restlessness | 0.6 ± 1.0 | 0.3 ± 0.6 | 0.1 |
| Performing repetitive mannerisms | 0.6 ± 1.2 | 0.4 ± 1.0 | 0.17 |

**Table 4:**
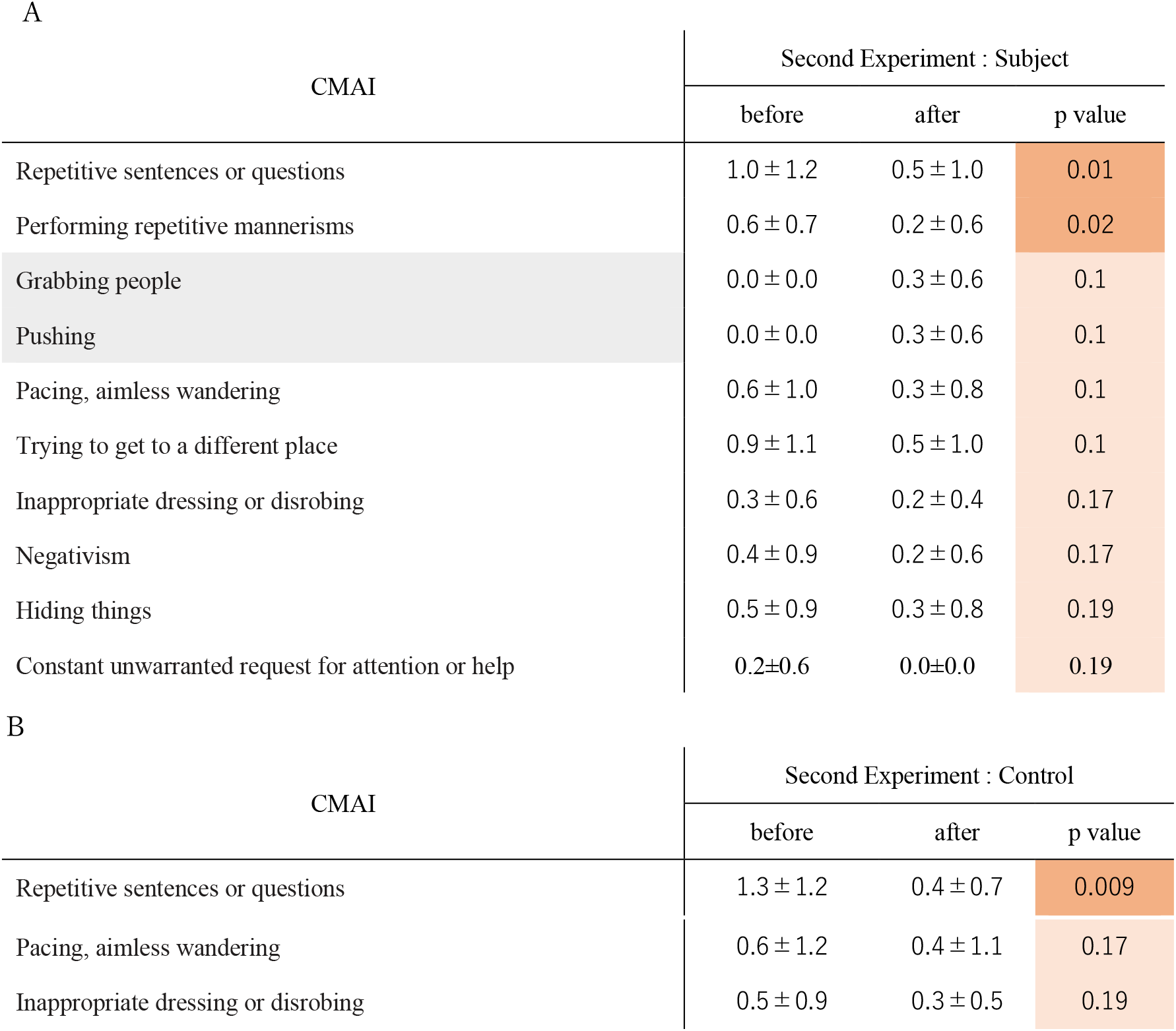
CMAI categories with trend and significant improvement in the Second experiment A: Subject, B: Control

In the second experiment improvement was significant in 5 categories in DBD: Shows lack of interest in daily activities (p=0.0004), Sleeps excessively during the day(p=0.002), Overeats(p=0.017), Wets himself/ herself (p=0.009), Soils himself/ herself (p=0.008), and 2 categories in Non-Aggressive CMAI; Repetitive mannerisms (p=0.01) and Repetitive sentences (p=0.008). Interestingly, the control group also improved in the Non-Aggressive CMAI significantly in the second experiment.

### 3. Memory test

Memory test scores were unchanged in both the first and second experiments. However, even though subjects did not improve their score, they responded to questions more willingly and some mentioned the season when they were asked “What day is today?” at the end of the 2^nd^ week.

## IV. Discussion

The most notable finding in this study was improvement in MMSE, DBD and CMAI during the experiment period, although no subject perceived the scent of eucalyptol. Table 5 shows the difference in MMSE, DBD, and CMAI scores before and after the two 1-week experiments for all subjects and the control group. “A” indicates the value of before the first experiment, “B” indicates after–before value of the first experiment, and “C” indicates after–before value of the second experiment. The positive number indicates improvement in MMSE and the negative number indicates improvement in DBD and CMAI. Blue indicates improvement, grey indicates the score before the 1^st^ experiment, and orange indicates residents who did not participate in any activity in each unit during the experiment week.

**Table 5.**
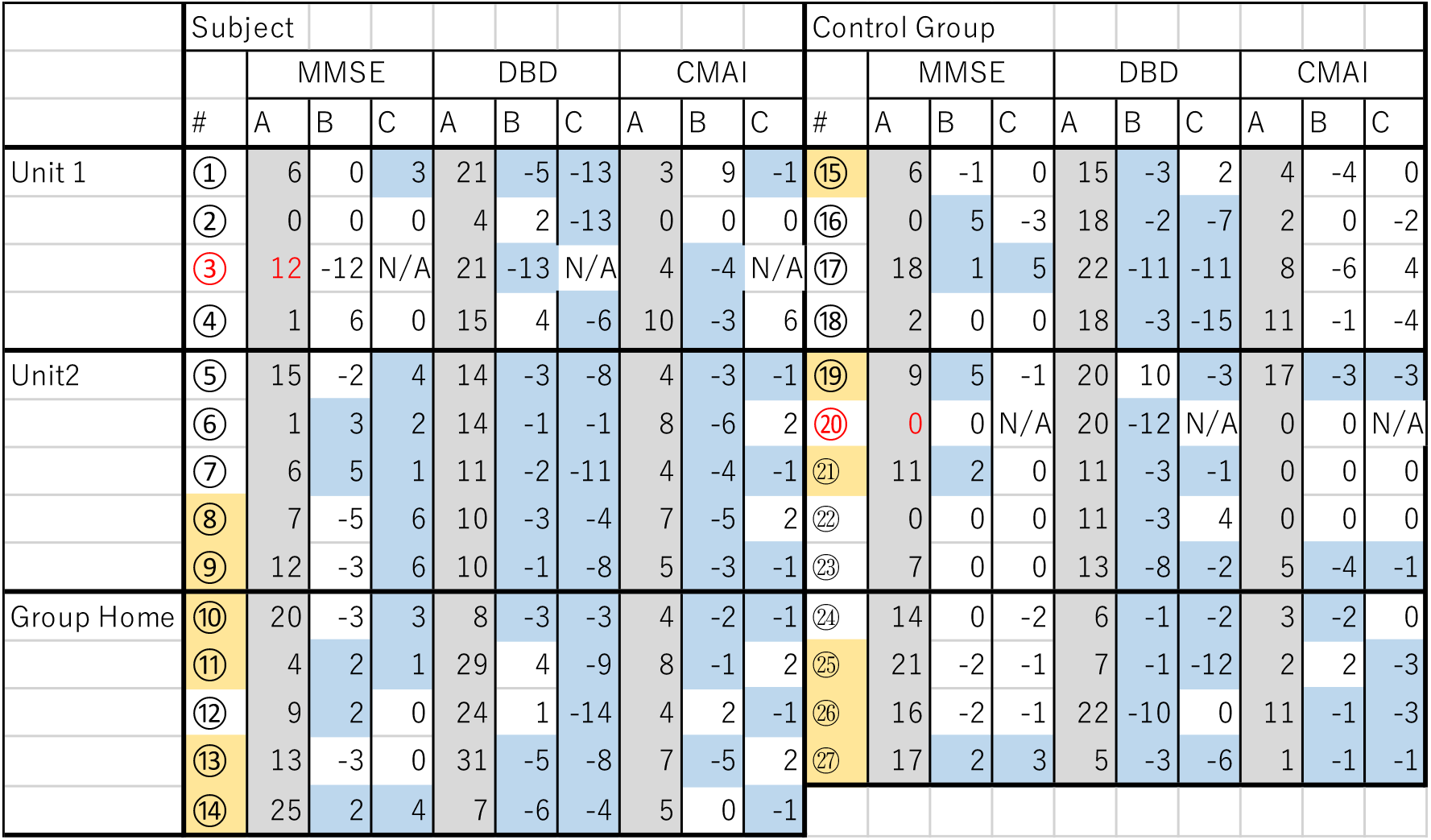
Changes in MMSE, DBD, and CMAI for the subject and control groups

The number of subjects who improved on the MMSE increased from two to all between the two experiments in Unit 2 in the 2^nd^ experiment (Fig.1). In Unit 2, the unit with the smallest private rooms and the common space among 3 units, all subjects improved their DBD score in both experiments, and all subjects improved their CMAI score in the first experiment. Among the three units, Unit 2 was the only unit where all subjects improved on the MMSE in the second experiment. Interestingly, there were residents who improved their MMSE, DBD, and CMAI during the experiments even in the control group, particularly in the first experiment. It is unexpected that the control group, which did not have the diffuser in their room, also showed the improvement only during the experiment week.

Although residents determine their own wake-up time in the facility, caregivers call them around their individual wake up time, help them change, and take them to the living room for a meal. As a result, all residents have breakfast and spend time in the living room after the meal while the doors to their rooms are kept wide open. Perhaps some diffused scent from the subject’s room entered the living room when the door was opened after the subjects woke, and both subjects and the control group experienced some effect from the scent drifting into the living room during the day. Because the additional scent in the second experiment was diffused 30 minutes before bed time and the door was closed while subjects were in their room, a little scent could have drifted from the subjects’ rooms. Perhaps, since the control group received little influence from the evening scent, their scores on the MMSE, DBD, and CMAI during the second experiment did not improve more than during the first experiment.

As Fig.1 shows, five units in the nursing home are divided by solid walls and their layouts differ. Among Unit 1, Unit 2, and the group home, the living room of Unit 2 is the smallest and located in front of the five subjects’ rooms. If the scent drifted to the living room, the concentration of scent in Unit 2 would have been the highest because of its size and location near the subjects’ room. Therefore, a big improvement of MMSE in Unit 2 in the second experiment might be associated with a high concentration of drifted eucalyptol scent.

Furthermore, although CMAI scores for aggressive behavior did not change significantly, on average, two subjects, one in Unit 1 and one in Unit 2, had their CMAI scores for aggression change for the worse in the second experiment.

Table 6 shows the MMSE and CMAI scores of these subjects. “A” indicates the value of the first experiment, “B” indicates value of the second experiment, and “C” indicates value of B-A. As the table shows, both subjects improved their MMSE and CMAI scores for aggressive behavior in the first experiment but these scores dropped between the two experiments. During the 2^nd^ experiment, although the subject in Unit 2 had improved MMSE and CMAI scores for non-aggressive behavior, but less than during the 1^st^ experiment, the subject in Unit 1 showed no improvement in either category in the second experiment. During the 2^nd^ experiment, for the subject of Unit 2, the caregiver reported that the subject woke up at midnight and had trouble going back to sleep, which was unusual. This may indicate that the concentration of eucalyptol scent was too strong for these two subjects, and the scent aggravated their aggressive behavior. Given that some residents in the control group had improved their MMSE, DBD and CMAI scores, probably just by being exposed to the drifting scent during the experiment weeks, the effect of the scent could differ depending upon the subtle amount of volatile components and their effect, which may differ by individual.

**Table 6.** MMSE and CMAI scores of the two subjects

|  |  | MMSE |  |  | CMAI Aggressive |  |  | CMAI Non-Aggressive |  |  |
| --- | --- | --- | --- | --- | --- | --- | --- | --- | --- | --- |
|  |  | A | B | C | A | B | C | A | B | C |
| Unit 1 | 1st | 1 | 7 | 6 | 8 | 5 | -3 | 2 | 2 | 0 |
|  | 2nd | 2 | 2 | 0 | 0 | 6 | 6 | 4 | 4 | 0 |
| Unit 2 | 1st | 1 | 4 | 3 | 2 | 0 | -2 | 6 | 2 | -4 |
|  | 2nd | 2 | 4 | 2 | 3 | 8 | 5 | 11 | 8 | -3 |

With respect to the effect of a subtle amount of volatile substance, Matsubara reported that a decline in attention during the visual discrimination task for 30 min. was significantly diminished by a low dose volatile of the leaves of *Laurus noblis L.*(laurel) though a high-dose volatile was not effective. ^20^ Because eucalyptol is the main component of *Laurus noblis L.*, this study may also confirm the effect of the drifting scent of eucalyptol. Although the effect of aromatherapy is often linked with a preference for the aroma,^21,22^ this study has shown that eucalyptol scent could influence MMSE, DBD and CMAI scores without olfactory perception by subjects. This means that eucalyptol was not only stimulating the olfactory cortex, but also the vomeronasal organ (VNO), the peripheral sensory organ of the accessory olfactory system which detects non-volatile chemical cues. Since neuronal signals from the VNO are sent to the hypothalamus, which affects behavior as well as other functions such as body temperature and perception of seasonal changes, this may explain how eucalyptol influenced DBD and CMAI scores.

Staff members in charge of each unit change shifts 3 times a day in this facility, and different caregivers take care of residents every morning. Therefore, it was impossible to get comments from caregivers based on continuous observation of residents. However, during the two experiments, caregivers commented on improvement of facial expression, better interaction, and decreased accidents as a result of subjects becoming more collaborative. All staff perceived that the eucalyptol scent in the subjects’ rooms masked other odors, so that staff could complete their tasks more comfortably given the fresher air. Thus, the eucalyptol scent not only improved the MMSE, DBD, and CMAI scores of subjects but also reduced the burden of caregivers.

The results of this pilot study indicate that eucalyptol has the potential to be an effective therapeutic option even at a level unperceivable to subjects; however, at the same time, it could be a factor in exacerbating aggressive behavior if too strong. Although this study provides evidence of the effect of eucalyptol on people with dementia, the number of subjects tested within one facility was very limited. Further research on the usage of eucalyptol at different durations and concentrations with subjects who have different types or severities of dementia is needed.

## Author Contribution

<u>Seiko Goto</u> supervised the study and wrote main manuscript <u>Hinako Suzuki</u> conducted the experiment and data analysis.

<u>Toshinori Nakagawa</u> and <u>Kuniyoshi Shimizu</u> tested the level of eucalyptol in the air of the experiment sites.

### Additional Information

There is no competing financial and non-financial interests.

